# Vascular and synaptic proteomes reveal blood-brain barrier disruption and postsynaptic remodeling in human temporal lobe epilepsy

**DOI:** 10.64898/2026.06.11.731740

**Authors:** Gavin Spillard, Mingzi Zhang, Nadia A. Atai, Allison Bosworth, Veronica A. Clementel, Jonathan J Russin, Charles Y. Liu, Marcelo P. Coba, Ruslan Rust

## Abstract

Blood-brain barrier (BBB) dysfunction and mesial temporal lobe epilepsy (MTLE) are considered to be engaged in a pathological feedback loop, with the consequent worsening of both conditions. However, the molecular landscape of the disruptions at the synapse and the blood-brain barrier during MTLE remains poorly characterized. Here, we perform quantitative proteomics on paired brain microvessel and vessel-depleted postsynaptic density (PSD) enriched fractions isolated from the epileptic hippocampus and ipsilateral temporal pole of patients with drug-resistant MTLE. The microvessel fraction (1,541 proteins; 439 differentially expressed proteins (DEPs)) reveals loss of tight-junction and endothelial adhesion proteins together with pericyte markers, concurrent with a significant increase of fibrinogen, plasminogen, complement C3, GFAP, and enrichment for complement and coagulation cascades. The PSD enriched fraction (7,450 proteins; 1,881 DEPs) shows the consequences of BBB leakage with an increase of protein infiltration, alongside inflammatory and extracellular-matrix proteins, together with disruption of the pre- and postsynaptic signaling machinery and loss of GABAergic interneurons. Cross-referencing healthy-brain expression confirms that the dysregulation of the processes reflects disease-associated changes rather than regional differences. Immunohistochemistry confirms microvascular remodeling, pericyte loss, parenchymal fibrinogen extravasation, microglial activation and presynaptic marker depletion in the epileptic hippocampus. Ligand-receptor mapping reveals dysregulation of the neurovascular ECM-adhesion interface, with upregulated parenchymal ECM ligands and downregulated vascular integrin receptors. Network-proximity analyses nominate candidate disease-modifying compounds for reversing the combined vascular and synaptic MTLE signature. Together, these findings establish a molecular map of vascular and synaptic dysfunction in human MTLE.

## Introduction

Epilepsy affects over 70 million people worldwide, yet one in three patients never achieves seizure freedom despite optimal pharmacotherapy.^1^ Mesial temporal lobe epilepsy (MTLE) is the most common form of drug-resistant focal epilepsy. A pathological hallmark of MTLE is hippocampal sclerosis, which is characterized by selective neuronal loss, reactive gliosis, and reorganization of synaptic circuits.^2–5^ Recent evidence from neuroimaging, cerebrospinal fluid biomarkers, and animal models suggests that blood-brain barrier (BBB) dysfunction and extravasation of blood-borne proteins into the brain parenchyma contributes to seizure generation and disease progression.^6–8^ For instance, albumin extravasation activates astrocytic TGF-beta signaling, and blood-derived proteins including fibrinogen promote neuroinflammation through microglial activation via CD11B and TLR4 receptors and complement deposition.^9–12^ In endothelial cells, the loss of Claudin 5 (CLDN5), a key tight junction protein that regulates blood brain barrier permeability, has been associated with increased seizure susceptibility and worse epilepsy outcomes.^8,13^ Moreover, BBB dysfunction not only contributes to the progression of seizures ^14,15^ but also leads to neuronal hyperexcitability and therefore can be considered a risk factor for MTLE.^16^ This produces a positive feedback loop that worsens the vasculature and neuronal conditions.^17^ However, the individual molecular components driving this vascular-synaptic crosstalk in human MTLE remain poorly characterized, masking disease-relevant signals confined to the vasculature or the synapse.

Prior proteomic studies of human MTLE tissue have relied on bulk homogenates^18^, therefore it was not possible to map the dysregulated molecular components of the vascular, neuronal and synaptic components. The postsynaptic density (PSD) contains a highly organized protein scaffold machinery at excitatory synapses, with the capacity to regulate a variety of synaptic functions through the regulation of different families of proteins including glutamate receptors, kinases, phosphatases GTPases and channels. While numerous studies have associated synaptic proteins to a variety of brain disorders, their direct characterization in human epilepsy remains poorly studied.^19^

To address this gap, we applied a dual-fraction proteomic approach. From surgically resected tissue of patients with drug-resistant MTLE, we isolated brain microvessels and a vessel-depleted PSD-enriched fraction from both the epileptic hippocampus and the ipsilateral temporal pole, enabling a paired within-patient comparison of vascular and synaptic proteomes. We complemented quantitative proteomic profiling with immunohistochemical validation and cross-referenced our results with healthy-brain expression data to distinguish disease-specific from region-specific changes. Ligand–receptor mapping revealed a dysregulated neurovascular ECM-adhesion interface, with upregulated parenchymal ECM ligands and downregulated vascular integrin receptors. Network-proximity analyses further nominated candidate disease-modifying compounds targeting the combined vascular and synaptic MTLE signature.

## Results

### Patients with temporal lobe epilepsy and study design

Patients with drug-resistant MTLE (4 female, 1 male; mean age 37.2 years, range 22–51; mean epilepsy duration 27.4 years) underwent resective surgery at two centers (Keck Hospital USC; Los Angeles General Medical Center). All had failed two or more antiepileptic drugs. Tissue was collected from the epileptic hippocampus and the ipsilateral temporal pole at surgery. MRI review classified the temporal pole as radiologically normal, showing no signal abnormalities or volume loss on T2-weighted and FLAIR sequences, while the hippocampus displayed hallmarks of mesial temporal sclerosis, including hippocampal atrophy and increased FLAIR signal intensity (**Fig. 1a,b**). From each tissue pair, we isolated brain microvessels and a vessel-depleted postsynaptic density (PSD) fraction enriched for synaptic scaffolding and signaling proteins (**Fig. 1b**). Additionally, matched tissue sections from the same patients were used for immunohistochemistry to validate key proteomic findings.

**Figure 1:**
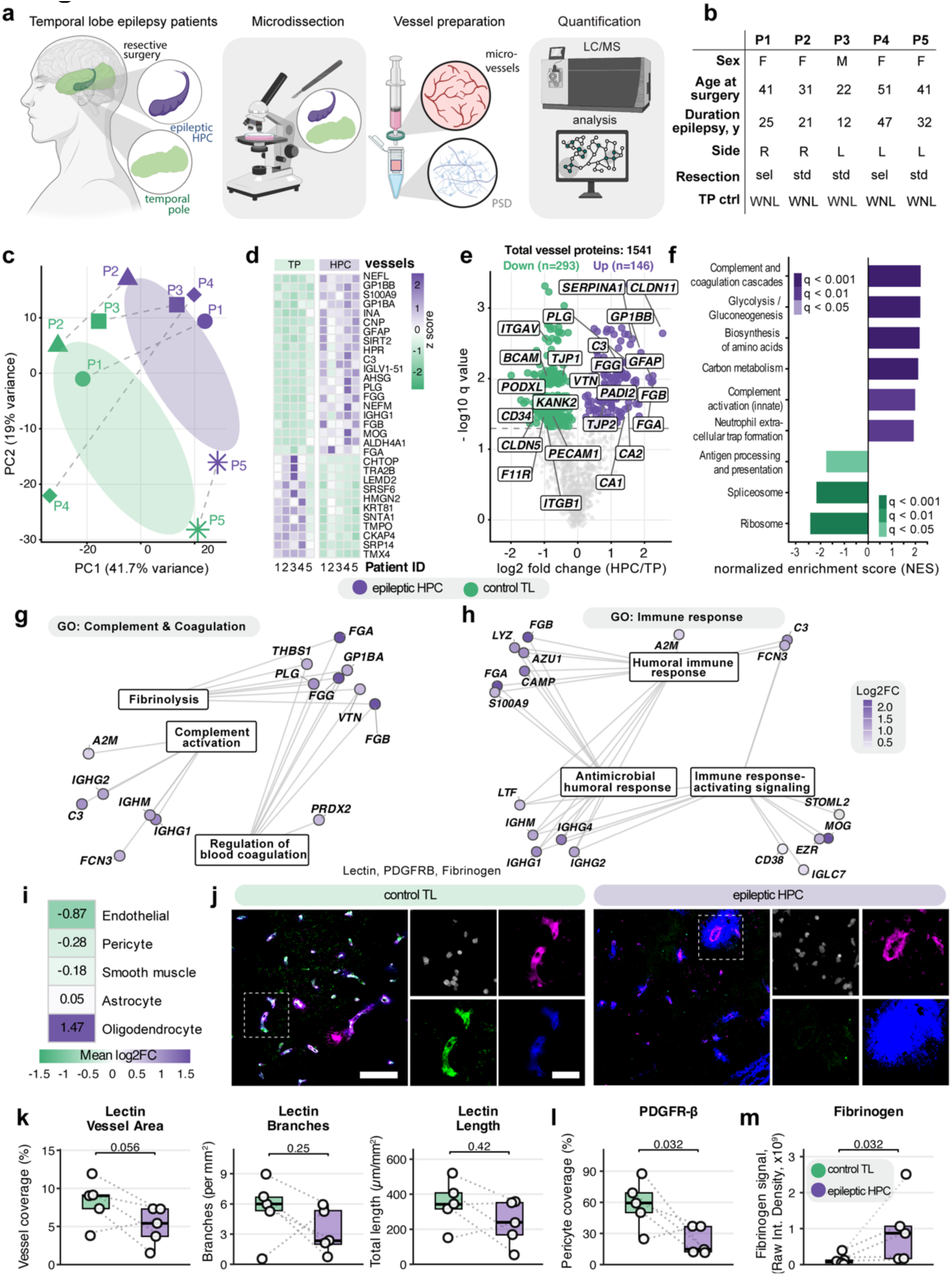
Study design and vascular compartment proteomics. **(a)** Schematic of the paired design: brain microvessels and vessel-depleted PSD-enriched fractions were isolated from the epileptic hippocampus and ipsilateral temporal pole of five patients with drug-resistant MTLE. **(b)** Clinical cohort: sex, age at surgery (years), epilepsy duration (years), resection side (L, left; R, right), resection type (std, standard anterior temporal lobectomy; sel, selective amygdalohippocampectomy) and MRI-based temporal pole status (WNL, within normal limits). **(c)** PCA of microvessel TMT intensities; dashed lines connect paired samples. **(d)** Heatmap of top DEPs across all five patients. **(e)** Volcano plot of microvessel DEPs (epileptic HPC vs. TP), selected proteins labelled (purple, higher in HPC; green, higher in TP). **(f)** KEGG enrichment of microvessel DEPs; bars, normalized enrichment scores; colour, adjusted significance. **(g)** GO network for complement and coagulation (fibrinolysis, complement activation, regulation of blood coagulation modules); node colour, log2FC. **(h)** GO network for immune response (humoral, antimicrobial humoral and immune response-activating signalling modules); node colour, log2FC. **(i)** Cell-type deconvolution of the microvessel fraction (endothelial, pericyte, smooth-muscle, astrocyte, oligodendrocyte); colour, mean log2FC. **(j)** Representative images from temporal pole and epileptic hippocampus stained for Lectin (vessels), PDGFRβ (pericytes) and Fibrinogen (extravasation); DAPI in blue. Scale bars, 50 µm (close-up, 20 µm). **(k)** Lectin-positive vessel area, branch number and total branch length per field (n = 5 pairs, paired t-test). **(l)** PDGFRβ-positive pericyte coverage of the vascular area (n = 5 pairs, paired t-test). **(m)** Parenchymal Fibrinogen immunoreactivity (raw integrated density; n = 5 pairs, paired t-test). Box plots show median with interquartile range; dots, individual patients; dotted lines connect paired samples.

### Epileptic hippocampal microvessels show BBB disruption and vascular inflammation

The High-Performance Liquid Chromatography Mass Spectrometry (HPLC-MS) analysis of brain microvessels were able to identify and quantitate 1,541 proteins. PCA analysis showed a clear separation between hippocampus and temporal pole along PC1 (41.7% variance), with paired samples from the same patient clustering together (**Fig. 1c**). We identified 439 DEPs (q ≤ 0.05), of which 146 (33%) were upregulated and 293 (67%) were downregulated in MTLE hippocampus (**Fig. 1d,e**).

Protein markers of vascular damage and neuroinflammation had a significant increase in their total protein levels. For instance, all three fibrinogen chains (FGA, FGB, FGG; log2FC +2.2-2.3) were among the top upregulated proteins, alongside, plasminogen (log2FC +1.39), complement C3 (log2FC +1.39), GFAP (log2FC +2.15), SERPINA1 (log2FC +1.65) PADI2 (log2FC +1.72) and platelet glycoprotein GP1BB (log2FC +1.89). Downregulated proteins in microvessels of the epileptic hippocampus suggested a major disruption of the BBB (**Suppl. Table 1**). HPLC-MS analysis shows a significant decrease CLDN5 (log2FC −1.61). CLDN5 is the most abundant claudin family member in endothelial cells from brain microvessels and it has been reported to be reduced in the hippocampus of MTLE patients when compared to postmortem sample analysis of controls.^8^ Moreover, it has been shown that inducible knockdown of claudin-5 in mice leads to spontaneous recurrent seizures.^8^

We also quantitated a significant decrease in cell adhesion and proteins that are essential for maintaining blood–brain barrier function during acute inflammation such as PECAM1, F11R, KANK2 TJP1, ITGAV, ITGB1, BCAM, and the PODXL family members CD34 and PODXL (**Fig. 1e and Suppl. Table 1**). Decrease in the protein levels of these molecules are not only associated to a disruption of the BBB, but also to a decrease in the capacity to regulate force transmission shear and the response to inflammation (**Fig. 1e and Suppl. Table 1**).^20–23^ This decrease in structural components of the BBB co-occur with an increase in the total protein levels of claudin 11 (CLDN11), TJP2, Carbonic anhydrase 1-2, and vitronectin, suggesting vascular and anti-inflammatory compensatory mechanisms (**Fig. 1e and Suppl. Table 1**).

KEGG pathway analysis revealed complement and coagulation cascades (q < 0.001), glycolysis/gluconeogenesis (q < 0.001), biosynthesis of amino acids (q < 0.001), and neutrophil extracellular trap formation (q ≤ 0.05) as significantly enriched among upregulated proteins, while downregulated pathways included carbon metabolism, spliceosome, and ribosome pathways, indicating impairment in metabolism and protein synthesis (**Fig. 1f**). GO enrichment network analysis revealed that nearly all differentially expressed proteins within fibrinolysis, complement activation, blood coagulation, and humoral immune response modules were upregulated in the epileptic hippocampus (**Fig. 1g,h**), indicating a coordinated activation of complement and coagulation cascades.

To better understand the cell-type specific contributions to the observed proteomic changes, we performed cell-type deconvolution using published brain scRNA-seq markers. We inferred shifts in the microvessel proteome composition, with decreased abundance of endothelial (mean log2FC = -0.87), pericyte (mean log2FC = -0.28), and smooth muscle cell marker proteins (mean log2FC = -0.18), and an enrichment of oligodendrocyte-derived proteins (mean log2FC = +1.47), possibly reflecting accumulation of myelin-derived material in the perivascular space (**Fig. 1i**).

To validate the MS analysis at the tissue level, we performed immunohistochemistry on matched hippocampal and temporal pole sections from the same patients (**Fig. 1j-m**). Lectin staining revealed a 38.8% reduction in vessel area in the epileptic hippocampus compared to the temporal pole (5.0 ± 2.5% vs. 8.3 ± 3.0%, p = 0.056, paired t-test), accompanied by trends toward reduced vascular branching and total vessel length (p = 0.25 and p = 0.42, respectively) (**Fig. 1j**). PDGFR-β immunolabeling showed a significant 61.4% decrease in pericyte coverage of microvessels (22 ± 13% vs. 58 ± 23%, p = 0.032; **Fig. 1k,l**), consistent with the downregulation of pericyte markers observed in the MS quantitative analysis. Extravascular fibrinogen immunoreactivity was 7.2-fold higher in the epileptic hippocampus (9.54 × 10⁸ ± 9.60 × 10⁸ vs. 1.32 × 10⁸ ± 1.51 × 10⁸ raw integrated density, p = 0.032; **Fig. 1m**), indicating substantial blood-brain barrier leakage.

In sum, these data provide convergent proteomic and histological evidence for microvascular rarefaction, pericyte loss, and BBB breakdown in the epileptic hippocampus, with specific loss of key tight-junction (CLDN5), adhesion (PECAM1, F11R) and pericyte (PDGFRB) proteins alongside compensatory upregulation of CLDN11 and TJP2, indicating a remodeling of the molecular landscape of the BBB.

### Synaptic proteome reveals neuroinflammation, pre/postsynaptic dysregulation, and mitochondrial dysfunction in the epileptic hippocampus

After isolating microvessels, we then prepared postsynaptic density (PSD) enriched fractions from each sample. This enrichment allowed us to obtain a high coverage of postsynaptic density proteins and a good coverage of presynaptic, dendritic and cell body proteins.^24–26^ With this pipeline, we were able to quantitate 7,450 proteins (**Fig. 2a**).

**Figure 2:**
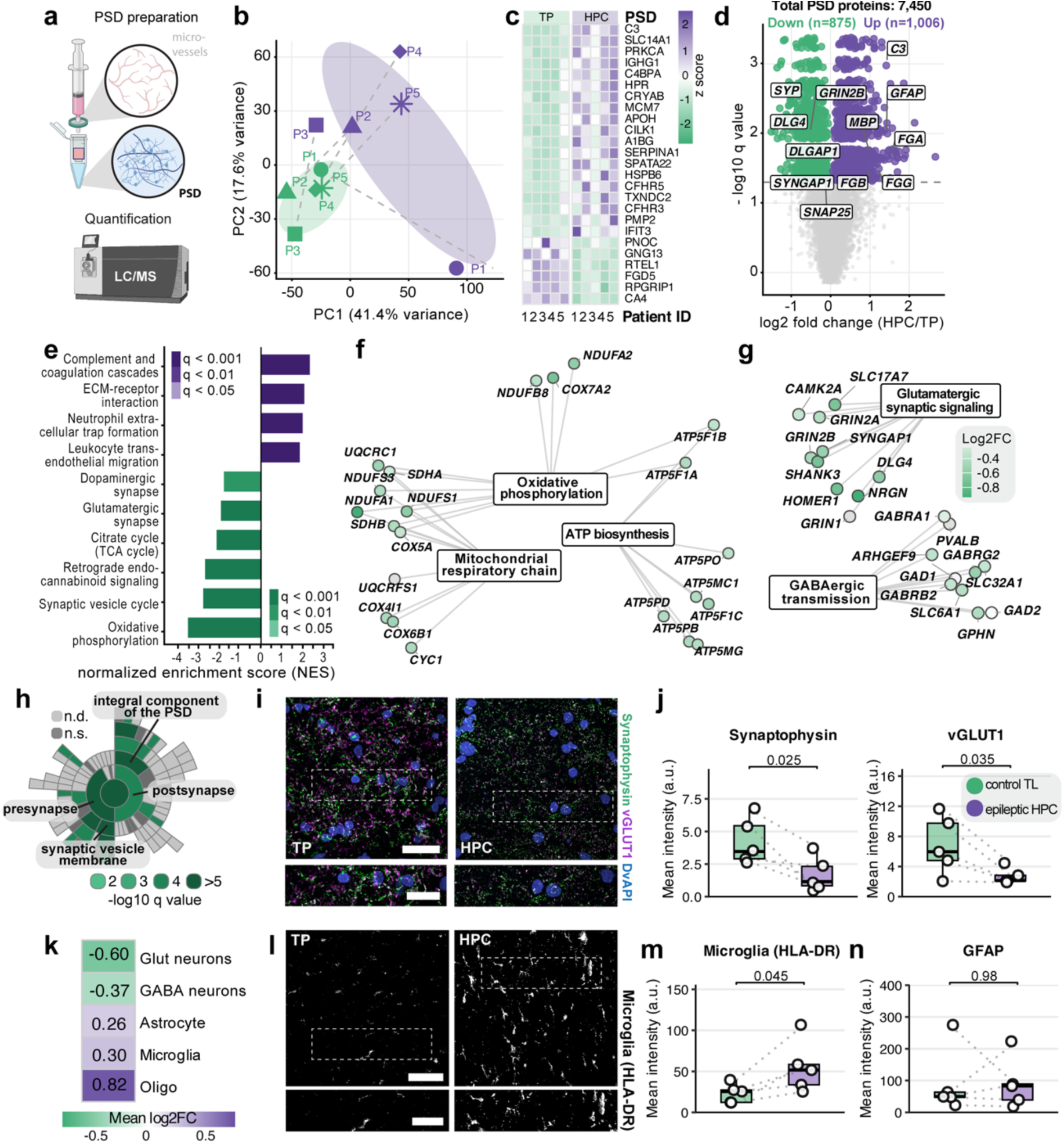
Synaptic compartment proteomics and immunohistochemical validation. **(a)** Workflow for isolation and proteomic analysis of the vessel-depleted PSD fraction. **(b)** PCA of the PSD fraction (PC1 = 41.4%, PC2 = 17.6% variance); dashed lines connect paired samples. **(c)** Heatmap of top DEPs across all five patient pairs (TP and HPC side by side). **(d)** Volcano plot of PSD DEPs (epileptic HPC vs. TP). **(e)** GSEA normalized enrichment scores for selected KEGG pathways; colour, adjusted significance. **(f)** PPI network of mitochondrial and oxidative phosphorylation DEPs (oxidative phosphorylation, mitochondrial respiratory chain, ATP biosynthesis modules); node colour, log2FC. **(g)** PPI network of synaptic DEPs (glutamatergic signalling and GABAergic transmission modules); node colour, log2FC. **(h)** SynGO sunburst of synaptic ontology terms enriched among PSD DEPs; colour, enrichment q-value (n.d., not defined; n.s., not significant). **(i)** Representative images of synaptic markers (Synaptophysin, vGLUT1; DAPI) in temporal pole and epileptic hippocampus. Scale bar, 20 µm. **(j)** Synaptophysin and vGLUT1 mean intensity (n = 5 pairs, paired t-test). **(k)** Cell-type deconvolution of the PSD fraction (glutamatergic neurons, GABAergic neurons, astrocytes, microglia, oligodendrocytes); colour, mean log2FC. **(l)** Representative images of microglia (HLA-DR) in temporal pole and epileptic hippocampus. Scale bar, 50 µm. **(m)** HLA-DR mean intensity (n = 5 pairs, paired t-test). **(n)** GFAP mean intensity (n = 5 pairs, paired t-test). Box plots show median with interquartile range; dots, individual patients; dotted lines connect paired samples.

PCA separated hippocampus from temporal pole (PC1 = 41.4% variance; **Fig. 2b**), and unsupervised hierarchical clustering of all differentially expressed proteins (DEPs) showed consistent within-group patterning across all five patient pairs (**Fig. 2c**). We identified 1,881 DEPs (q ≤ 0.05), of which 1,006 (53%) were upregulated and 875 (47%) downregulated in the MTLE hippocampus (**Fig. 2d**). Among the most significantly downregulated proteins were key components of the pre- and postsynaptic signaling machinery of excitatory synapses, including synaptophysin (SYP), DLG4, GRIN2B, DLGAP1, SYNGAP1, SHANK3, and SNAP25, where as complement C3, GFAP, fibrinogen chains (FGA, FGB, FGG), and myelin basic protein (MBP) were among the top upregulated hits in the epileptic hippocampus (**Fig. 2d**).

KEGG pathway analysis revealed oxidative phosphorylation as the most strongly suppressed pathway (NES = −3.52, q < 0.001), followed by synaptic vesicle cycle (NES = −2.73, q < 0.001), retrograde endocannabinoid signaling (NES = −2.72, q < 0.001), citrate cycle (q < 0.01), and glutamatergic synapse (q ≤ 0.05; **Fig. 2e**). Among upregulated KEGG pathways, complement and coagulation cascades (NES = +2.36, q < 0.001), neutrophil extracellular trap formation (q < 0.001), leukocyte transendothelial migration (q < 0.001), and ECM-receptor interaction (q < 0.01) were the top hits (**Fig. 2e**), suggesting a significant effect of vascular dysfunction in the MTLE hippocampus.

Consistent with oxidative phosphorylation as the most strongly suppressed pathway, GO enrichment network analysis resolved these mitochondrial DEPs into three interconnected modules: oxidative phosphorylation, mitochondrial respiratory chain, and ATP biosynthesis (Fig. 2g). Notably, we map this deficit to specific protein complexes, including NADH dehydrogenases (NDUFA1, NDUFS1, NDUFB8), cytochrome c oxidase subunits (COX5A, COX6B1), and ATP synthase components (ATP5F1A, ATP5PO) (**Fig. 2f and Suppl. Table 1**), extending previous reports of mitochondrial dysfunction in MTLE hippocampus.^27–29^

GO enrichment network analysis further resolved these downregulated synaptic DEPs into two connected modules: a glutamatergic module comprising core PSD scaffold and signaling proteins (CAMK2A, GRIN2A/B, DLG4, SYNGAP1, SHANK3, HOMER1, SLC17A7), and a GABAergic module anchored by GAD1, GAD2, and GABRB2, with markers of somatostatin- (SST), pro-neuropeptide Y (NPY), and cholecystokinin-positive (CCK) interneurons also reduced (**Fig. 2g**).

SynGO analysis of the downregulated PSD proteins revealed synaptic impairment in both pre-and postsynaptic compartments, with the presynaptic compartment more prominently affected. The most enriched terms were integral component of synaptic vesicle membrane (4.9-fold enrichment, q = 4.2 × 10^−10^; 23 genes including SYP, SYT1, SV2A/B), synaptic vesicle proton loading (6.4-fold, q = 4.0 × 10^−7^), and synaptic vesicle endocytosis (3.0-fold, q = 6.1 × 10^−4^), whereas postsynaptic density membrane components were also affected but at lower enrichment (3.0-fold, q = 4.0 × 10^−7^; 30 genes including GRIA1-4, GRIN2B, and LRRTM2-4; **Fig. 2h**). Consistent with this synaptic loss signature in HPLC-MS, immunohistochemical quantification showed that synaptophysin immunoreactivity was reduced by 60% (paired t-test, p = 0.025) and vGLUT1 by 61% (p = 0.035) in the epileptic hippocampus relative to the temporal pole (**Fig. 2i,j**).

Cell-type deconvolution inferred shifts in the PSD fraction composition, with decreased abundance of glutamatergic neuron markers (mean log2FC = -0.60; including CAMK2A, DLG4, GRIN2A, and SLC17A7) and GABAergic interneuron markers (mean log2FC = -0.37; **Fig. 2k**), and a concomitant increase of glial markers: microglia (mean log2FC = +0.30; AIF1, CSF1R, CX3CR1), astrocytes (mean log2FC = +0.26; GFAP, AQP4, S100B), and oligodendrocytes (mean log2FC = +0.82; MBP, PLP1, MOG, **Fig. 2k**). Together, these data suggest a neuroinflammatory phenotype in the epileptic PSD fraction, characterized by microglial activation and reactive astrogliosis alongside neuronal and synaptic loss.

Immunohistochemistry confirmed increased microglia activation in the epileptic hippocampus, with HLA-DR immunoreactivity increased by 137% relative to the control temporal pole (paired t-test, p = 0.045; **Fig. 2l,m**). Notably, GFAP immunoreactivity did not differ significantly between regions (p = 0.98; **Fig. 2n**), which likely reflects the different sensitivities of HPLC-MS in biochemically enriched fractions versus IHC in brain tissue sections.

Together, these data demonstrate that the epileptic hippocampus undergoes broad synaptic remodeling with pre- and postsynaptic disruption of core signaling machineries, accompanied by mitochondrial dysfunction, neuroinflammation with activated microglia.

### Disease-specific and region-specific proteomic changes in the epileptic hippocampus

Because our paired design compares the epileptic hippocampus with the ipsilateral temporal pole, observed differences could in principle be affected by normal regional biology rather than disease. To separate these contributions, we cross-referenced all DEPs with GTEx v8 brain expression data (regions: brain hippocampus vs. brain cortex) and classified each protein as disease-specific (|log2FC| ≤ 1 in GTEx), region-specific (same direction with |log2FC| > 1.0 in GTEx), disease-amplified (same direction but stronger in epilepsy), or disease-opposed (opposite direction; **Fig. 3a,b**).

**Figure 3:**
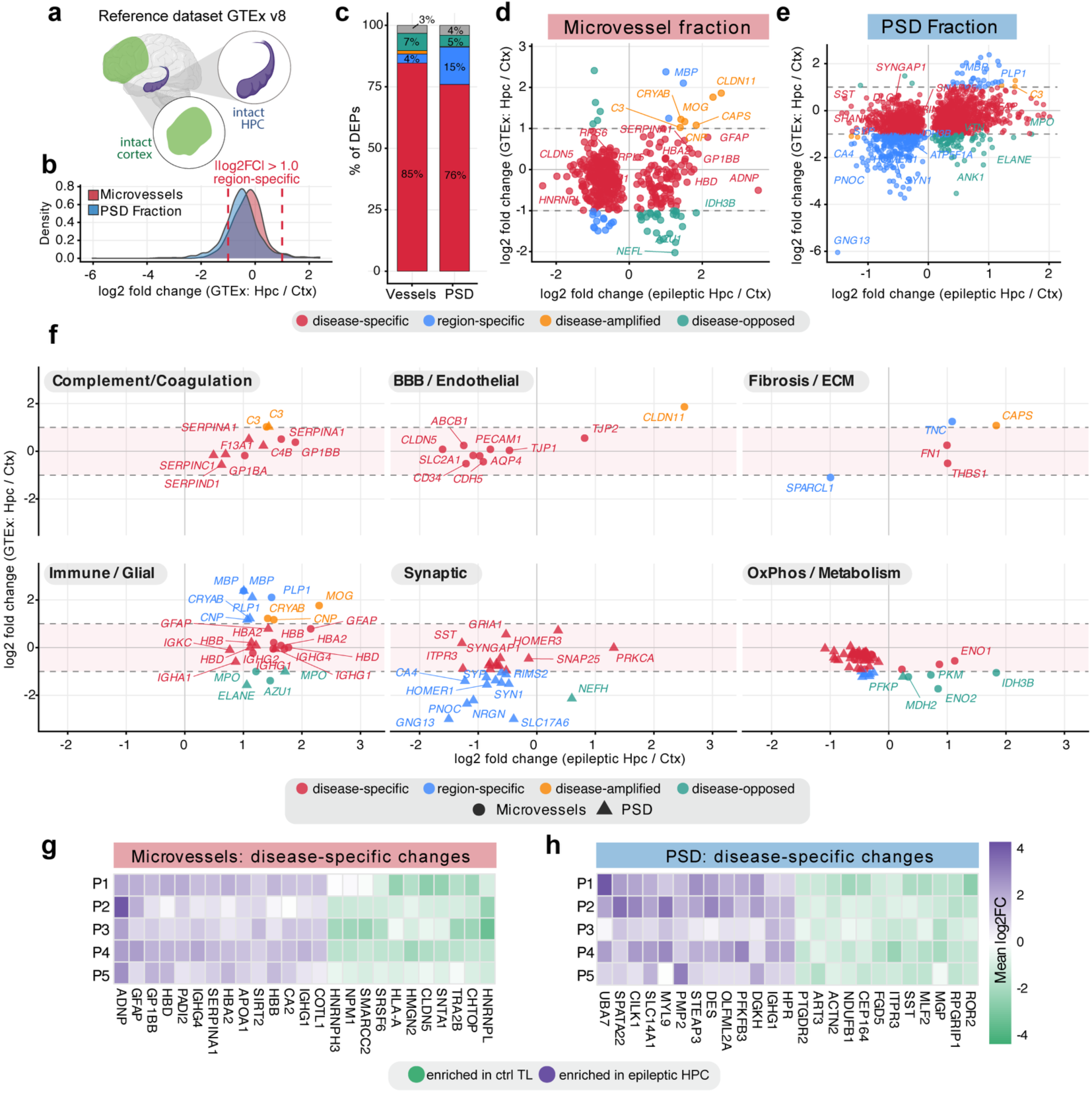
Disease-specific versus region-specific proteomic changes. **(a)** Schematic of the GTEx v8 healthy-brain cross-reference classifying each DEP as disease-specific, region-specific, disease-amplified or disease-opposed. **(b)** Density distribution of GTEx Brain_Hippocampus vs. Brain_Cortex log2FC for all quantified microvessel (red) and PSD (blue) proteins; dashed line, the |log2FC| > 1.0 threshold flagging baseline regional differences. **(c)** Proportion of DEPs in each class for the microvessel and PSD fractions. **(d,e)** Joint scatter of epileptic HPC/TP log2FC against GTEx HPC/Ctx log2FC for the microvessel (d) and PSD (e) fractions, coloured by class. **(f)** Same decomposition stratified by six functional themes (complement/coagulation, BBB/endothelial, fibrosis/ECM, immune/glial, synaptic, OxPhos/metabolism). Circles, microvessel DEPs; triangles, PSD DEPs. **(g,h)** Per-patient heatmaps of disease-specific DEPs in the microvessel (g) and PSD (h) fractions across the five pairs.

Across both fractions, the majority of DEPs were classified as disease-specific (microvessels, ∼85%; PSD, ∼76%), with a smaller region-specific component that was more pronounced in the PSD fraction (PSD: ∼15% vs microvessels: 4%), consistent with the larger baseline transcriptional divergence between hippocampus and cortex at the neuronal level (**Fig. 3c**). Joint scatter plots of the epileptic HPC/TP log2FC against the GTEx HPC/Ctx log2FC visualized this decomposition at single-protein resolution for both the microvessel (**Fig. 3d**) and PSD (**Fig. 3e**) fractions. For instance, the downregulated tight-junction and endothelial adhesion proteins CLDN5 and PECAM1 fell into the disease-specific quadrant of the microvessel scatter (**Fig. 3d**), while myelin-and stress-related proteins such as CLDN11, MOG and CRYAB clustered among the disease-amplified hits. In the PSD scatter the downregulated interneuron marker SST, together with synaptic regulators such as SYNGAP1 and ANK1, fell into the disease-specific cluster (**Fig. 3e**). To interpret the disease-specific signal along biological themes, we stratified the scatter by six functional categories from relevant KEGG pathways (**Fig. 3f**). In the complement and coagulation panel, fibrinogen chains (FGA, FGB, FGG) together with SERPINA1/A3 and GP1BB were largely disease-specific in both the microvessel and PSD fractions. The BBB/endothelial panel confirmed CLDN5, PECAM1 and ABCB1 loss as disease-specific and the upregulation of CLDN11 as disease-amplified. The fibrotic/ECM and immune/glial signature was likewise disease-specifically upregulated in both fractions, with the matrix/fibrosis components FN1 and THBS1 and the astroglial marker GFAP were elevated alongside neutrophil granule proteins (MPO, ELANE, AZU1) and immunoglobulin components (IGHG1–4, IGHA1, IGKC, HBA2, HBB, HBD), consistent with vascular fibrosis, reactive gliosis, granulocyte infiltration, and plasma/erythrocyte leakage across a compromised BBB. In contrast, the myelin/oligodendrocyte proteins (MBP, PLP1, CNP) segregated as region-specific rather than disease-specific, reflecting baseline white-matter differences between hippocampus and temporal pole. Interestingly, in the synaptic panel loss of SYP, SYT1, SV2A, GRIA1–4, SNCA and HOMER3 was largely disease-specific, although a small subset of synaptic markers (e.g., NRGN, SLC17A6, HOMER1) was classified as region-specific, suggesting that hippocampal–cortical differences might also play a role in the observed quantitative differences in the MTLE hippocampus. Notably, the OxPhos/metabolism panel revealed in the PSD fraction the broad depletion of NDUFA/NDUFS, SDHA and ATP5 subunits was essentially entirely disease-specific, whereas in the microvessel fraction several glycolytic and TCA-cycle enzymes (PKM, IDH3B, MDH2, ENO2) fell into the disease-opposed quadrant; these enzymes are normally enriched in hippocampus relative to cortex in healthy brain, suggesting that the epileptic hippocampal vasculature has lost its normal metabolic profile (**Fig. 3f**). Additionally, per-patient heatmaps of the top disease-specific DEP set (GTEx |log2FC| ≤ 1.0) (**Fig. 3g,h**) showed that these pathological signatures were consistent across all five patients. Together these findings confirm that the identified differences in BBB impairment, fibrinogen upregulation, synaptic loss and mitochondrial dysfunction, reflect epileptic pathology rather than baseline regional differences.

### Dysregulated ligand-receptor signaling in hippocampal MTLE

To resolve inter-compartment signaling disrupted in MTLE, we mapped DEPs from both fractions onto curated ligand–receptor (target) pairs from the OmniPath database (6,658 validated human pairs) and retained a curated set of 53 GTEx-filtered cross-fraction pairs after excluding baseline region-specific changes (**Fig. 4a**). We separated these pairs into two directions: (1) vessel-to-PSD pairs, in which microvascular ligands engage synaptic receptors, and (2) PSD-to-vessel pairs, in which synaptic or parenchymal ligands engage vascular receptors.

**Figure 4:**
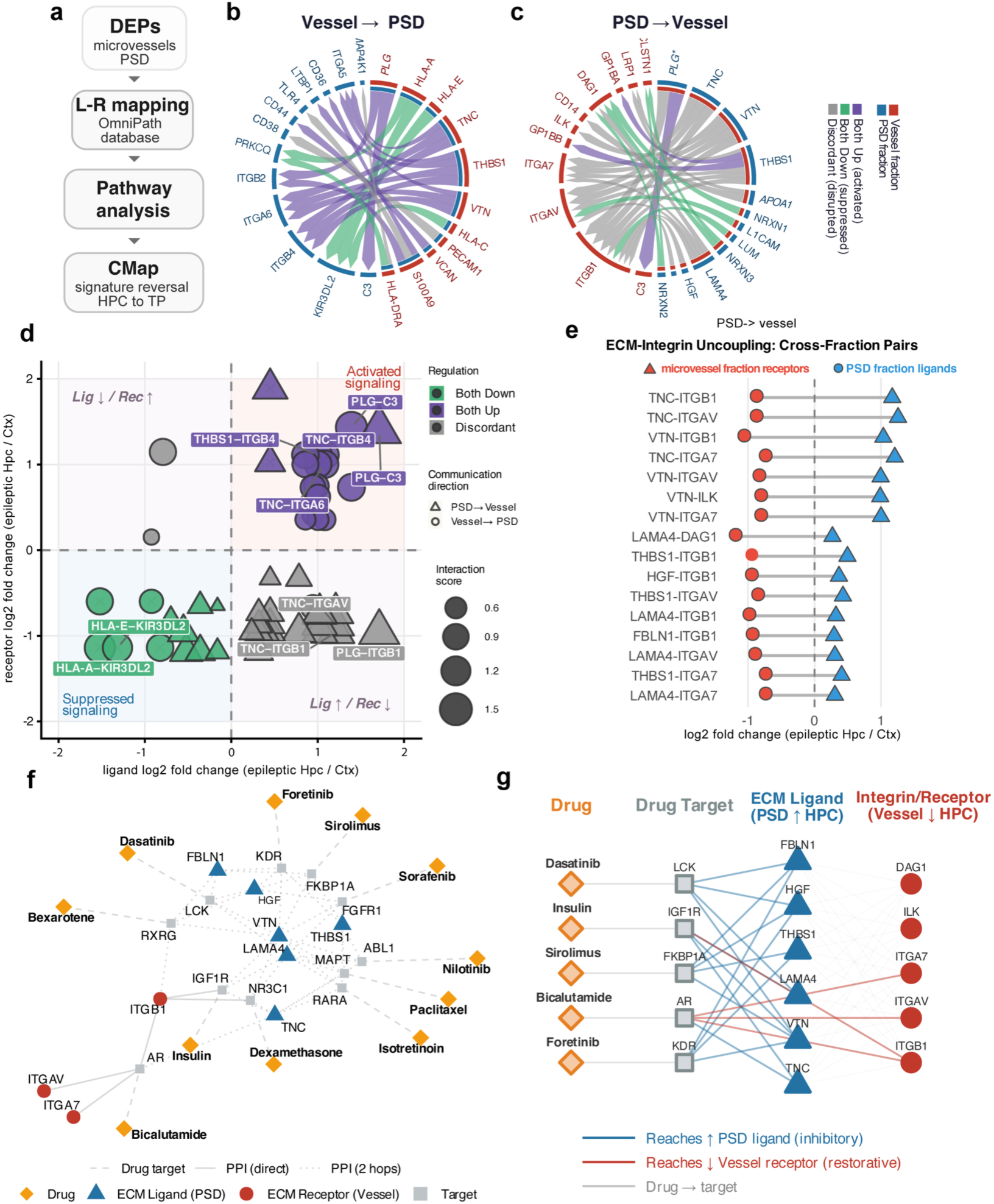
Dysregulated ligand–receptor interactome and in silico drug repurposing. **(a)** Workflow: disease-specific DEPs from both fractions were mapped onto the OmniPath ligand–receptor database, analysed for pathway enrichment, and used to query the Connectivity Map (CMap) L1000 reference for HPC-to-TP signature reversal. **(b)** Chord diagram of vessel-to-PSD pairs, with microvascular ligands (red outer arc) engaging PSD-side receptors (blue outer arc). Chord colour, regulation class (purple, both up/activated; green, both down/suppressed; grey, discordant). **(c)** Reciprocal PSD-to-vessel pairs, with parenchymal/synaptic ligands (blue outer arc) engaging vascular receptors (red outer arc); same colour code. **(d)** Ligand–receptor fold-change scatter: ligand log2FC (x) and receptor log2FC (y) for each pair. Shape, communication direction (circles, vessel→PSD; triangles, PSD→vessel); colour, regulation class; size, interaction score (mean |log2FC|). Quadrants annotate activated, suppressed and discordant regulation. **(e)** Cross-fraction ECM–integrin pairs (PSD-to-vessel) ranked by opposing regulation. Blue triangles, PSD ECM ligand log2FC; red circles, microvessel receptor log2FC. **(f)** PPI network linking CMap compounds (orange diamonds) through their targets (grey squares) to ECM ligands (blue triangles) and vascular integrin receptors (red circles). Dashed lines, drug–target; solid lines, direct PPIs; dotted lines, two-hop PPIs. **(g)** Simplified hierarchical view showing each drug, its target, and whether it reaches an upregulated ECM ligand (blue, inhibitory) or a downregulated vascular receptor (red, restorative).

In the vessel-to-PSD direction, upregulated vascular ligands (PLG, TNC, THBS1, VTN, VCAN, S100A9) engaged PSD receptors spanning integrin (ITGA6, ITGB2), complement (C3), and innate-immune (CD14, TLR4, CD36) (**Fig. 4b**). In the PSD-to-vessel direction, ECM ligands (TNC, VTN, THBS1, LAMA4, HGF, LUM) engaged vascular integrins (ITGB1, ITGAV, ITGA7), ILK, and LRP1; neurexins (NRXN1, NRXN2, NRXN3) paired with DAG1 and CLSTN1, canonical synaptic adhesion interactions; and L1CAM paired with ITGAV (**Fig. 4c**). A ligand–receptor fold-change scatter showed that most pairs signaling from vessel-to-PSD fell into the activated-signaling quadrant, with e.g., PLG–C3, TNC–ITGB4, TNC–ITGA6, and PLG–ITGB2 among the most strongly co-activated (**Fig. 4d**). Interestingly, we found most discordant ligand–receptor pairs clustered in the PSD-to-vessel direction, predominantly showing upregulation of parenchymal ligands and downregulation of their cognate vascular receptors (**Fig. 4d**). Most of these discordant pairs mapped to ECM–integrin interactions (**Fig. 4e**). ECM ligands in the PSD fraction (TNC, VTN, THBS1, LAMA4, HGF, FBLN1) were consistently upregulated in the epileptic hippocampus, whereas their cognate integrin receptors in the vessel fraction (ITGB1, ITGAV, ITGA7), together with integrin-linked kinase ILK and dystroglycan DAG1, were downregulated. This opposing regulation suggests a direct effect of increased parenchymal signaling on the vascular system, with the consequent uncoupling of the neurovascular ECM–adhesion interface, characterized by increased parenchymal matrix components alongside reduced vascular receptor availability.

To identify compounds predicted to reverse the epileptic proteomic signature, we performed Connectivity Map (CMap) analysis on the vessel, PSD, and combined signatures (**Fig. 4f,g**). The top-ranked hits spanned three drug classes: kinase inhibitors (foretinib, sorafenib, vemurafenib, sirolimus), nuclear receptor ligands (isotretinoin, bexarotene, bicalutamide), and hormone/metabolic modulators (insulin, anastrozole) (**Fig. 4f,g)**. To position these candidates relative to the dysregulated neurovascular interface, we layered the CMap hits onto a protein–protein interaction network and computed shortest paths to the ECM–integrin module. Foretinib, sirolimus, sorafenib, bexarotene, and isotretinoin sat closest to ECM ligands and vascular integrin receptors (**Fig. 4f,g**), pointing to two complementary modes of action: suppression of elevated parenchymal signals and restoration of reduced receptor pathways, although these computational predictions require experimental validation. For example, foretinib, through inhibition of KDR, modulates downstream signaling from HGF and THBS1, while bicalutamide, via AR, may restore ITGB1 and ITGAV (**Fig. 4g**).

Together, these analyses identify a convergent signaling network associated with the disrupted ECM–integrin signaling axis linking vascular and synaptic compartments in MTLE.

## Discussion

Our simultaneous profiling of the vascular and synaptic compartments from the same patients reveals that specific components of the coagulation and complement cascades drive a pathological inflammatory feedback loop between the BBB and the synapse in the epileptic human hippocampus. Fibrinogen chains, plasminogen, and complement C3 emerge as key mediators of vascular-to-neuronal inflammatory signaling. In parallel, the neurovascular ECM-adhesion interactions show a pattern of uncoupling, potentially driven by increased parenchymal matrix deposition that reduces vascular receptor availability.

Fibrinogen chains and plasminogen were among the most consistently upregulated proteins across both fractions, and plasminogen-C3 interactions were the top cross-compartment ligand-receptor interactions. Fibrinogen and plasminogen are not expressed in normal brain parenchyma, making their presence in both the microvessel and PSD fractions a marker of BBB leakage. Previous work in multiple sclerosis and Alzheimer’s disease models established that extravasated fibrinogen activates microglia through CD11b/CD18 and promotes complement deposition.^9,12^ Our data extend this paradigm to human MTLE and suggest that fibrinogen-mediated neuroinflammation links BBB disruption to synaptic dysfunction. Similarly, plasminogen is a major component of neuroinflammatory processes at the BBB. Plasminogen is converted into plasmin, which increases BBB permeability, promotes immune cell infiltration, and facilitates extracellular matrix degradation.^30,31^ It triggers inflammation by stimulating microglial activation, generating inflammatory cytokines, and disrupting tight junctions, often involving pericyte-mediated signaling. Accordingly, we observed disruption of key tight junction proteins including CLDN5, which has been previously linked to MTLE^8,13^ and also compensatory tight-junction remodeling of the CLDNs and TJP proteins in the microvessel fraction, consistent with compensatory tight-junction remodeling reported in neurological and psychiatric conditions with BBB compromise.^32^

Notably, the detection of high levels of fibrinogen and plasminogen in the PSD-enriched fraction indicates that blood-derived proteins penetrate deep into the synaptic compartment, potentially interfering directly with postsynaptic signaling.^33^ This inflammatory cascade may be further exacerbated by the observed pericyte loss. The proteomic and histological evidence for pericyte depletion (PDGFR-β reduction of 61.4%, p = 0.032) is consistent with prior reports of pericyte depletion in experimental MTLE models and human hippocampal sclerosis.^34–36^ Pericytes actively regulate BBB permeability, cerebral blood flow, and immune cell trafficking, and their loss may therefore amplify BBB dysfunction beyond what tight-junction disruption (as observed for e.g. CLDN5) alone would produce.^37–39^ Whether pericyte loss is a cause or consequence of seizure activity remains unresolved, though experimental evidence that pericyte depletion itself can trigger spontaneous seizures^40^ supports a bidirectional relationship.

The suppression of oxidative phosphorylation in the PSD fraction (NES = −3.52) represents one of the strongest pathway signals reported in MTLE proteomics. This aligns with evidence that synaptic mitochondria are selectively vulnerable in neurodegeneration and suggests that energy failure at the synapse is a primary pathological feature of MTLE. The concurrent loss of synaptic vesicle cycle proteins implies that impaired mitochondrial ATP production directly compromises neurotransmitter release.^28^ Here, we observe that specific families of proteins interconnected in oxidative phosphorylation, mitochondrial respiratory chain, and ATP biosynthesis processes were particularly disrupted. This includes genes involved in Mendelian disorders of mitochondrial function such as NDUFA1 (mitochondrial complex I deficiency, nuclear type 12), NDUFS1 (mitochondrial complex I deficiency, nuclear type 5), NDUFB8 (mitochondrial complex I deficiency, nuclear type 32), COX5A (mitochondrial complex IV deficiency, nuclear type 20), COX6B1 (mitochondrial complex IV deficiency, nuclear type 7), ATP5F1A (mitochondrial complex V deficiency, nuclear type 4A/B), and ATP5PO (mitochondrial complex V deficiency, nuclear type 7).

The opposing regulation of parenchymal ECM ligands (e.g., TNC, VTN, THBS1) and their cognate vascular integrin receptors (e.g., ITGB1, ITGAV) represents a new feature of the epileptic neurovascular unit. ECM-integrin signaling is essential for maintaining vascular integrity and BBB permeability.^41,42^ The upregulation of ECM ligands in the parenchymal compartment likely reflects reactive matrix remodeling in response to seizure activity^43^, while the concurrent loss of vascular integrin receptors may impair endothelial and pericyte responsiveness to these signals, further destabilizing the BBB. This uncoupling pattern is consistent with ECM/integrin pathway dysregulation identified in transcriptomic studies of epilepsy models^44^ and with the role of integrin signaling in seizure susceptibility.^45^

The paired within-patient design controls for inter-individual variability and represents a key methodological strength. Nevertheless, the cohort size (n = 5), while typical for studies using resected human brain tissue, limits statistical power for detecting modest effect sizes or stratifying by clinical variables. The ipsilateral temporal pole serves as the best available within-patient control, though it may be influenced by chronic seizure activity; GTEx cross-referencing provides an independent healthy-brain reference to mitigate this concern. The CMap drug repurposing predictions offer a computational starting point but are derived from cancer cell line signatures rather than BBB-specific models and require future experimental validation. Isolation of microvessels from human tissue can also retain a degree of adjacent white matter, which may contribute to the myelin-associated signal in this fraction; our region-specific decomposition (Fig. 3f) helps separate this component from the disease-driven vascular changes. Finally, the cross-sectional nature of the tissue precludes establishing temporal causality between BBB disruption and synaptic changes; longitudinal or animal model studies, together with functional permeability assays, will be needed to address directionality.

In conclusion, our dual-fraction proteomic approach reveals that BBB disruption and synaptic degeneration in the epileptic hippocampus are molecularly connected through shared inflammatory cascades and ECM–integrin uncoupling. These findings establish a molecular map of vascular and synaptic dysfunction in human MTLE and nominate specific protein interactions and candidate compounds as targets for future therapeutic investigation. Because the two compartments are molecularly linked, targeting them together rather than either alone may prove therapeutically advantageous.

## Methods

### Patient selection and recruitment

Patients with drug-resistant temporal lobe epilepsy (TLE) were identified through a prospectively maintained clinical database at Keck Hospital of USC and Los Angeles General Medical Center (formerly LAC+USC Medical Center). Eligibility was determined using predefined criteria to ensure a clinically well-characterized cohort. Inclusion criteria were: (i) medically refractory TLE meeting the International League Against Epilepsy (ILAE) definition of drug resistance and requiring surgical intervention; (ii) availability of comprehensive preoperative clinical data and neuroimaging; and (iii) documentation of perioperative and postoperative outcomes. Exclusion criteria were incomplete medical records, insufficient imaging for analysis, and severe pre-existing neurological deficits unrelated to epilepsy that could confound interpretation. All cases were reviewed by a multidisciplinary epilepsy surgery board prior to resection.

Five patients met these criteria (four female, one male; age at surgery 22–51 years, mean 37.2 ± 11.1 years; age at seizure onset 4–16 years, mean 9.8 ± 4.3 years; epilepsy duration 12–47 years, mean 27.4 ± 13.1 years). Surgical procedures comprised selective amygdalohippocampectomy (n = 2; P1, P4) or standard anterior temporal lobectomy (n = 3; P2, P3, P5), performed on the right (n = 2; P1, P2) or left (n = 3; P3, P4, P5) hemisphere based on presurgical evaluation. Diagnosis of mesial temporal sclerosis (MTS) was made on preoperative 3T MRI, where all five epileptic hippocampi displayed hippocampal atrophy and increased T2/FLAIR signal intensity, and was confirmed post-operatively by neuropathological examination of the resected hippocampus by a board-certified neuropathologist according to ILAE classification (Type 1, gliosis only, or no MTS). The ipsilateral temporal pole was within normal limits. Clinical and demographic variables are summarized in Fig. 1b. Both sexes were included; sex-based analyses were not performed given the limited sample size, which represents a limitation to generalizability.

The study was conducted in accordance with the Health Insurance Portability and Accountability Act and approved by the University of Southern California Institutional Review Board (protocol HS-20-00038). All participants provided written informed consent for surgical treatment, tissue donation, and the collection of clinical and demographic data prior to tissue collection.

### Tissue collection

Specimens were obtained from patients undergoing en bloc resection as part of standard surgical treatment for medically refractory TLE, with procurement adhering to strict ethical and institutional guidelines. For each patient, two tissue specimens were collected intraoperatively: the epileptic hippocampus (disease tissue) and the ipsilateral temporal pole (within-patient control). Tissue was placed on ice immediately upon resection and processed in parallel without delay to preserve molecular integrity, for microvessel isolation and postsynaptic density (PSD) fractionation. Prior to homogenization, visible white matter was removed from each tissue block under a dissecting microscope, and white matter was further depleted during the dextran density-gradient step

### Brain microvessel isolation and PSD fractionation

PSD enriched samples were isolated after the removal of temporal pole or hippocampus microvessels. Both samples were prepared as we described before^24,25,46–48^. Fraction purity using this protocol has been established previously, ^24,25,46–48^ and was confirmed here by the enrichment of compartment-specific markers in the proteomic data.

Microvessels samples were further washed by addition of 1mL ice cold PBS/vortex mixed/centrifuged at 1,200 rpm at 4^0^C for 10 min and then removal of PBS (x3). Both regions from each patient were homogenized and washed in parallel under identical conditions, and microvessels were washed three times in ice-cold PBS to remove residual luminal blood prior to lysis, controlling for intravascular blood as a source of plasma-protein signal. Samples were kept on ice during the entire process. After the final washing step, we added 100uL lysis buffer (0.5M TEAB, 0.05% SDS). Samples were subjected to tip sonication (Q700, QSonica, amplitude = 10, 2 sec on/2 sec off pulses, 20 sec total processing time per sample, on ice) and then centrifuged at 15K rpm at 4^0^C for 10 min. The supernatant was then transferred to a fresh tube. The protein concentration of each sample was measured using the Qubit Protein Assay Kit (Thermo, Q33211) and Qubit 4.0 fluorometer per manufacturer’s instructions. Equal amount of protein (10 µg) per sample was transferred to a fresh tube adjusted to highest volume (90 µL) with lysis buffer. Four µL Reducing Reagent (Sigma, 4381664) were added to each sample. Samples were incubated at 60^0^C for 1hr. Two µL Alkylating Reagent (Sigma, 4381664) were added to each sample. Samples were incubated at room temperature for 15 min. Four µg trypsin/LysC (Promega, V50703) were added to each sample. Samples were incubated overnight at room temperature in dark.

TMTpro reagents (Thermo, A44522) were equilibrated at room temperature. Twenty µL of anhydrous acetonitrile (Sigma, 900644) were added to each label and the contents were then transferred to each sample. Samples were then incubated at room temperature for 1hr. Eight µL 5% hydroxylamine were added to each sample. Samples were incubated at room temperature for 15 min. Samples were then combined and dried-up using a speedvac (Eppendorf 5301 vacufuge concentrator). Peptides were mixed and subjected to offline alkaline C8 reverse phase fractionation. Twenty fractions were collected in a time-dependent manner and dried-up using a speedvac. These were desalted using C18 tips with 100 µL bed per manufacturer’s instructions (Thermo Scientific Pierce, 87784).

PSD samples were subjected to tip sonication (Q700, QSonica, amplitude = 10, 2 sec on/2 sec off pulses, 20 sec total processing time per sample, on ice) and then centrifuged at 15K rpm at 4^0^C for 10 min. The supernatant was transferred to a fresh tube (on ice) and the protein concentration of each sample was measured using the Qubit Protein Assay Kit (Thermo, Q33211) and Qubit 4.0 fluorometer per manufacturer’s instructions. Equal amount of protein (100 µg) per sample was transferred to a fresh tube adjusted to highest volume (90 µL) with lysis buffer. Four µL Reducing Reagent (Sigma, 4381664) were added to each sample. Samples were incubated at 60^0^C for 1hr. Two µL Alkylating Reagent (Sigma, 4381664) were added to each sample. Samples were incubated at room temperature for 15 min. Four µg trypsin/LysC (Promega, V50703) were added to each sample. Samples were incubated overnight at room temperature in dark.

TMTpro reagents (Thermo, A44522) were equilibrated at room temperature. Twenty µL of anhydrous acetonitrile (Sigma, 900644) were added to each label and the contents were then transferred to each sample. Samples were then incubated at room temperature for 1hr. Eight µL 5% hydroxylamine were added to each sample. Samples were incubated at room temperature for 15 min. Samples were then combined and dried-up using a speedvac (Eppendorf 5301 vacufuge concentrator).

To remove DOC, labelled peptides were combined in 1% TFA solution, vortex mixed and centrifuged at 15k rpm for 10 min. The supernatant was transferred to a fresh tube and dried-up using a speedvac (Eppendorf 5301 vacufuge concentrator). Peptides were then subjected to offline alkaline C8 reverse phase fractionation. Twenty fractions were collected in a time-dependent manner and dried-up using a speedvac. These were desalted using C18 tips with 100 µL bed per manufacturer’s instructions (Thermo Scientific Pierce, 87784).

### High-Performance Liquid Chromatography Mass Spectrometry (HPLC-MS)

The desalted peptide fractions of projects A and B were reconstituted in water 0.1% formic acid and analyzed using LC-MS with FAIMS ion mobility pre-separation (nano-easy LC 1200, Thermo Orbitrap Exploris 480). Raw data files of the LC-MS analysis were submitted to Proteome Discoverer 2.5 (Thermo) separately for target decoy search using Sequest against the homo sapiens canonical swissprot database (TaxID= 9606, v2021-10-30). The search allowed for up to two missed cleavages, a precursor mass tolerance of 20ppm, a minimum peptide length of six and a maximum of three equal dynamic modifications of oxidation (M), deamidation (N, Q) or phosphorylation (S, T, Y). Methylthio (C) and TMTpro (K, peptide N-terminus) were set as static modifications. Peptide level confidence was set at q ≤0.05. Only unique peptides were considered for protein quantitation.

Normalization mode of total peptide amount was applied. All mass spectrometry proteomics data have been deposited to the ProteomeXchange Consortium via the PRIDE partner repository.

### Bioinformatic analyses

All computational analyses were performed in R (v4.4.2) using RStudio using previously established workflows.^49–55^

Quality control and principal component analysis. All quantified proteins were subjected to principal component analysis (PCA) after median-centring each protein across samples and imputing remaining missing values with the row mean. Proteins with zero variance were excluded. Data were centered and scaled (unit variance) prior to PCA using the prcomp function. Sample-to-sample Pearson correlation matrices were computed on the full quantified proteome using pairwise complete observations; all within-fraction correlations exceeded r = 0.90. PCA and correlation plots were generated to assess data quality and confirm separation of hippocampal and temporal pole samples

#### Gene set enrichment analysis (GSEA)

GSEA was conducted on the full quantified proteome to detect coordinated pathway-level changes. All proteins were ranked by their mean log2FC in decreasing order (positive values indicating upregulation in epileptic hippocampus). Gene symbols were converted to Entrez IDs; duplicate entries were removed by retaining the first occurrence. GSEA was performed using clusterProfiler against three gene set collections: KEGG pathways (gseKEGG, organism “hsa”), GO Biological Process (gseGO), and MSigDB Hallmark gene sets (v2023.2; GSEA function with TERM2GENE). Parameters were set as: minimum gene set size = 10, maximum gene set size = 500, p-value cutoff = 0.05, p-value adjustment method = Benjamini-Hochberg, and random seed = 42 for reproducibility. Normalized enrichment scores (NES) were used to rank and visualize pathway-level changes.

#### Cell-type deconvolution

Cell-type deconvolution was performed similar to previous work.^47^ To estimate cell-type composition changes between epileptic hippocampus and temporal pole, we scored the mean log₂FC of detected marker proteins using curated marker gene sets for eight brain cell types: endothelial cells (23 markers), pericytes (14 markers including PDGFRB, RGS5, ABCC9, MCAM), astrocytes (15 markers including GFAP, AQP4, ALDH1L1), oligodendrocytes (15 markers including MBP, MOG, PLP1), microglia (14 markers including CX3CR1, P2RY12, TMEM119), excitatory neurons (14 markers including SLC17A7, CAMK2A, DLG4), inhibitory neurons (15 markers including GAD1, GAD2, SST, PVALB), and smooth muscle cells (12 markers including ACTA2, MYH11, CNN1). Marker gene sets were compiled from published single-cell RNA-seq references and the Allen Brain Atlas. For each cell type, we computed three metrics across the full quantified proteome (not restricted to DEPs): the number of detected marker proteins, the mean log₂FC of detected markers, and the percentage of markers upregulated in the epileptic hippocampus.

#### Protein-protein interaction (PPI) network analysis

PPI networks were constructed using the STRINGdb R package (v11.5, species code 9606 for Homo sapiens). For each fraction, the top DEPs ranked by absolute log2FC were mapped to STRING identifiers; unmapped proteins were excluded. Interactions were retained at a combined confidence score ≥ 400 (medium-high confidence). Undirected networks were constructed using the igraph package, and isolated nodes (degree = 0) were removed. Hub proteins were identified as the top 20 nodes ranked by degree centrality. For visualization, the top nodes by degree were displayed using the Fruchterman-Reingold layout algorithm (via ggraph and tidygraph), with node color encoding log2FC and edge width proportional to the STRING combined score.

#### Ligand-receptor interaction analysis

To map intercellular signaling between the vascular and synaptic compartments, we performed a cross-fraction ligand-receptor (L-R) analysis. A curated L-R interaction database was compiled by integrating pairs from OmniPath (ligrecextra resource, 6,658 human pairs), CellChat, and NicheNet, supplemented by manual annotation for brain-relevant interactions. The initial set comprised 100 ligand families spanning vascular growth factors (VEGFA/B/C, ANGPT1/2, PDGF family, FGF1/2), TGF-β superfamily (TGFB1–3, BMP2/4/7), Wnt signaling (WNT3A/5A/7A), chemokines and cytokines (CXCL12, CCL2, IL-6, TNF), neurotrophic factors (BDNF, NGF, GDNF), semaphorins, ephrins, neurexins, complement and coagulation factors (C3, C4A, PLG, SERPINE1), ECM proteins (COL4A1/2, FN1, LAMA1/2, THBS1/2, TNC), and neuropeptides (SST, NPY, VIP). L-R pairs were evaluated in two communication directions: vessel-to-PSD (microvascular ligands engaging PSD-side receptors) and PSD-to-vessel (parenchymal/synaptic ligands engaging vascular receptors). Pairs were retained only when both the ligand and receptor were quantified and had non-missing fold-change values. An initial set of 64 pairs (28 vessel-to-PSD, 36 PSD-to-vessel) was subjected to manual curation to remove pairs not relevant to the neurovascular unit, pairs where neither protein was detected, and biologically implausible interactions, yielding a final curated set of 53 L-R pairs. Each pair was classified by concordance (both proteins regulated in the same direction versus discordant regulation) and assigned to one of eight functional pathway groups: ECM-integrin, synaptic adhesion, complement/coagulation, vascular signaling, kinase signaling, lipid/metabolic, immune-vascular, and other. Chord diagrams were generated using the circlize R package.

*SynGO* SynGO analysis was performed using the SynGO web platform (https://www.syngoportal.org, release 20231201). Background was set to all detected proteins quantified in the PSD fraction. Enrichment was reported as fold enrichment with Benjamini–Hochberg–adjusted q-values; terms with q ≤ 0.05 were considered significant.

#### Cross-fraction integration

Proteins identified as differentially expressed in both the microvessel and PSD fractions were identified by intersection analysis. For overlapping DEPs, fold-change concordance was assessed by comparing the direction and magnitude of log2FC between fractions. DEPs were classified as concordant (same direction in both fractions: up-up or down-down) or discordant (opposite directions). The top 40 overlapping proteins ranked by combined magnitude (|log2FCmicrovessels| + |log2FCPSD|) were visualized using ComplexHeatmap.

#### Disease-specificity analysis using GTEx

To distinguish disease-driven proteomic changes from baseline regional differences between hippocampus and cortex, we cross-referenced our DEPs with healthy-brain transcriptomic data from the Genotype-Tissue Expression (GTEx) project (v8). Median transcript-per-million (TPM) values for Brain_Hippocampus and Brain_Cortex were obtained from the GTEx Portal (https://gtexportal.org). For each DEP, the corresponding GTEx log2FC (hippocampus/cortex) was computed. Proteins with |GTEx log2FC| > 1.0 were flagged as exhibiting substantial baseline regional expression differences. DEPs were then classified into four categories: disease-specific (significant change in epilepsy, minimal baseline regional difference), region-specific (change mirrors normal regional differences), disease-amplified (epilepsy change in the same direction as but exceeding the baseline regional difference), and disease-opposed (epilepsy change opposite to the baseline regional difference).

#### In silico drug repurposing via the Connectivity Map

Disease-specific DEPs from both fractions were used to query the Connectivity Map (CMap) L1000 reference dataset to identify compounds whose transcriptional signatures oppose the epilepsy-associated proteomic signature (hippocampus-to-temporal pole reversal). Candidate compounds were linked to their primary drug targets, and a protein-protein interaction network was constructed to connect CMap hits through their targets to dysregulated ECM ligands and vascular integrin receptors identified in the L-R analysis. Drug-target-DEP connectivity was assessed via direct PPIs and two-hop interactions in the STRING network. The disease-specific DEP signature (upregulated and downregulated gene lists) was submitted to the Connectivity Map (CMap) L1000 CLUE platform (clue.io, Broad Institute) using the ’touchstone’ compound dataset. Reversal scores were computed as the mean of the tau scores for the upregulated and downregulated query gene sets, with positive scores indicating signature reversal. Compounds were ranked by reversal score and filtered for significance (p ≤ 0.05 after permutation testing). Drug class annotations were obtained from the CLUE compound metadata.

### Immunohistochemistry and image analysis

Matched frozen sections (10 µm) from the epileptic hippocampus and temporal pole of all five patients were processed for immunofluorescence staining. After blocking with 10% normal donkey serum for 1 hour at room temperature, sections were incubated overnight at 4 °C with the following primary antibodies: anti-fibrinogen (Dako, 1:200), anti-PDGFRβ (R&D Systems, 1:20), DyLight 594-conjugated Lycopersicon esculentum lectin (Vector Laboratories, 1:200) for vessel labeling, anti-GFAP (Millipore Sigma, 1:1000), anti-HLA-DR/DP/DQ (Novus Bio, clone CR3/43, 1:50) for microglia/macrophage activation, anti-synaptophysin (Cell Signaling, 1:200), and anti-vGLUT1 (Cell Signaling, 1:200). Species-appropriate Alexa Fluor-conjugated secondary antibodies (Invitrogen, 1:200) were applied for 1 hour at room temperature, and nuclei were counterstained with DAPI. For the vascular integrity panel, triple-labeling was performed with lectin, anti-PDGFRβ, and anti-fibrinogen on the same sections.

Images were acquired on a confocal laser scanning microscope at 10x or 20x magnification, with identical acquisition parameters (laser power, gain, pinhole size) maintained across all samples within each staining panel. For each patient and brain region, 2–3 non-overlapping fields of view were captured. All image quantification was performed in ImageJ/FIJI (v1.54) using standardized, blinded analysis protocols. For lectin staining, images were binarized using consistent thresholds and vessel area (percent positive area), branch number, and total branch length were quantified. Fibrinogen extravasation was quantified as the raw integrated density of extravascular fibrinogen signal (parenchymal signal outside the lectin-positive vascular area). Pericyte coverage was calculated as the percentage of lectin-positive vessel area covered by PDGFRβ-positive signal. For synaptic markers (synaptophysin, vGLUT1), mean fluorescence intensity was measured after binarization at consistent thresholds. For microglial (HLA-DR) and astrocyte (GFAP) markers, mean fluorescence intensity was quantified across the full field of view.

### Statistical analysis

For each patient, the protein abundance ratio between the epileptic hippocampus and the ipsilateral temporal pole (HPC/TP) was computed and log2-transformed, such that positive values indicate higher abundance in the epileptic hippocampus. Differential abundance was determined in Proteome Discoverer 2.5 from these within-patient hippocampus/temporal pole ratios (n = 5 paired patients). Multiple testing correction was performed using Storey’s q-value method; proteins with q ≤ 0.05 were considered differentially expressed. For all pathway enrichment analyses (ORA and GSEA), p-values were corrected using the Benjamini-Hochberg procedure. For immunohistochemistry quantification, values from multiple fields per region were averaged within each patient to yield one value per patient per condition. Statistical comparisons between temporal pole and epileptic hippocampus were performed using two-tailed paired t-tests (n = 5 pairs), which account for the within-patient design. Data are presented as box plots showing median and interquartile range, with individual patient data points overlaid and connected by dashed lines to indicate pairing. All IHC statistical analyses were performed using the rstatix and ggpubr R packages.

## Acknowledgments

We thank the patients and their families for consenting to tissue donation, and the Keck Hospital of USC and Los Angeles General Medical Center (formerly LAC+USC Medical Center) neurosurgery teams for sample collection.

## Author Contributions

M.P.C. and R.R. designed the study. J.J.R., C.Y.L. and N.A. performed the surgical resections and provided clinical tissue samples. G.S., M.Z. and A.B. processed the brain tissue and prepared microvessel and PSD subcellular fractions. M.P.C., V.A.C. and R.R. performed the HPLC-MS and proteomic analyses. G.S., M.Z., R.R. analysed the immunohistochemical data. M.P.C. and R.R. generated the figures and wrote the manuscript. M.P.C. and R.R. supervised the project. All authors discussed the results and approved the final version of the manuscript.

## Data Availability

The mass spectrometry proteomics data have been deposited to the ProteomeXchange Consortium with the dataset identifier PXD077336. GTEx v8 data were accessed from the GTEx Portal (https://gtexportal.org) on February, 2026. An interactive data explorer is available at https://rustlab1.github.io/bbb-proteome-explorer/.

